# RNA degradation analysis reveals ribosome dynamics in complex microbiome samples

**DOI:** 10.1101/2021.04.08.439066

**Authors:** Susanne Huch, Lilit Nersisyan, Maria Ropat, Donal Barret, Jing Wang, Jaime Huerta-Cepas, Wu Wei, Lars M Steinmetz, Lars Engstrand, Vicent Pelechano

## Abstract

Post-transcriptional regulation is essential for life, yet we are currently unable to investigate its role in complex microbiome samples. Here we discover that co-translational mRNA degradation, where the degradation machinery follows the last translating ribosome, is conserved across prokaryotes. By investigating 5’P mRNA decay intermediates, we obtain *in vivo* ribosome protection information that allows the study of codon and gene specific ribosome stalling in response to stress and drug treatment at single nucleotide resolution. We use this approach to investigate *in vivo* species-specific ribosome footprints of clinical and environmental microbiomes and show for the first time that ribosome protection patterns can be used to phenotype microbiome perturbations. Our work paves the way for the study of the metatranslatome, and enables the investigation of fast, species-specific, post-transcriptional responses to environmental and chemical perturbations in unculturable microbial communities.

## Main Text

The microbiome is a key player in health and disease(*1, 2*). Its interaction with the host can lead to differential drug sensitivity or be associated with disease states(*1, 3*). Thus, it is necessary to understand how complex microbial communities respond to environmental changes and interact with the host. Metagenomics investigates microbiome composition by DNA content, while metatranscriptomics measures mRNA abundances in complex microbial mixtures(*4, 5*). However, neither approach informs about post-transcriptional regulation, which is essential to understand how the produced mRNA translates into diverse phenotypes(*6*). Metaproteomics is a useful approach, but can only provide a limited view of species-specific proteomes(*7*), as most identified peptides are shared across species. Furthermore, approaches like ribosome profiling, which have been key to dissect ribosome dynamics(*8*), cannot be applied to complex microbiome communities. Thus, there is an urgent need for approaches allowing for *in vivo* measurement of ribosome footprints in complex microbiomes.

We have previously demonstrated that studying co-translational mRNA degradation in yeast provides *in vivo* ribosome dynamics information(*9*). Co-translational mRNA degradation, where the eukaryotic 5’-3’ exonuclease Xrn1 follows the last translating ribosome is widespread in eukaryotic organisms, such as yeast(*9*) and plants(*10–12*). However, in prokaryotes, mRNA degradation was primarily thought to initiate via endonucleolytic cleavage followed by 3’-5’ degradation(*13–15*). More recent studies have demonstrated that 5’-3’ RNA exonucleolytic activity can also occur in prokaryotes. For example, the ribonuclease J was identified in *Bacillus subtilis*(*16*) and homologues are present in other species(*17*). However, the direct interaction between the translation process and mRNA degradation has not been investigated experimentally across the prokaryotic tree of life. Thus, we do not know yet if co-translational degradation significantly contributes to shaping the prokaryotic degradome. And if it does, how conserved is this process across prokaryotes and which degradation pathways are responsible.

To answer these questions, we investigate 5’ monophosphorylated mRNA decay intermediates (5’P) in isolated species and microbial communities. We show that co-translational mRNA degradation is common among prokaryotes and that this process involves multiple RNA degradation pathways. Finally, we demonstrate that 5’P RNA molecules serve as a good readout for *in vivo* ribosome dynamics in clinical and environmental microbiomes, thus opening an avenue for study of the metatranslatome.

## Results

### 5’-3’ co-translational mRNA degradation is common in prokaryotes

To investigate the potential existence of co-translational decay in prokaryotes(*9, 18*), we studied the 5’P mRNA degradome in cultured and complex bacterial communities using optimized 5PSeq(*9, 19–21*) (Table S1 and S2). We first investigated the 5’P degradome in open reading frames (ORFs) in *B. subtilis* (Fig. 1A). This revealed a clear 3-nucleotide (nt) periodicity of 5’P counts (Fig. 1B) as previously described for yeast(*9*), with a clear 5’P preference for the second nucleotide of each codon (F1), both at the metagene and single gene levels (Fig. 1C and Fig. S1A). 5’P degradation intermediates accumulate 11 and 14 nucleotides upstream of translation start and stop sites respectively, as expected from slow initiating and terminating ribosomes, analogous to the −14 nt from start and −17 nt from stop observed in budding yeast(*9, 21*). The smaller protection size can easily be explained by the known difference in ribosome size between eukaryotes and prokaryotes(*22*). To confirm the association of 5’P decay intermediates with translating ribosomes, we performed 5PSeq on polyribosomal fractions of sucrose density gradients and observed similar patterns (Fig. S1B). Finally, to demonstrate its biological origin, we confirmed that *in vitro* fragmentation of the same RNA nearly eliminated the observed *in vivo* 3-nt periodicity and start/stop associated footprints (Fig. 1A).

**Fig. 1.**
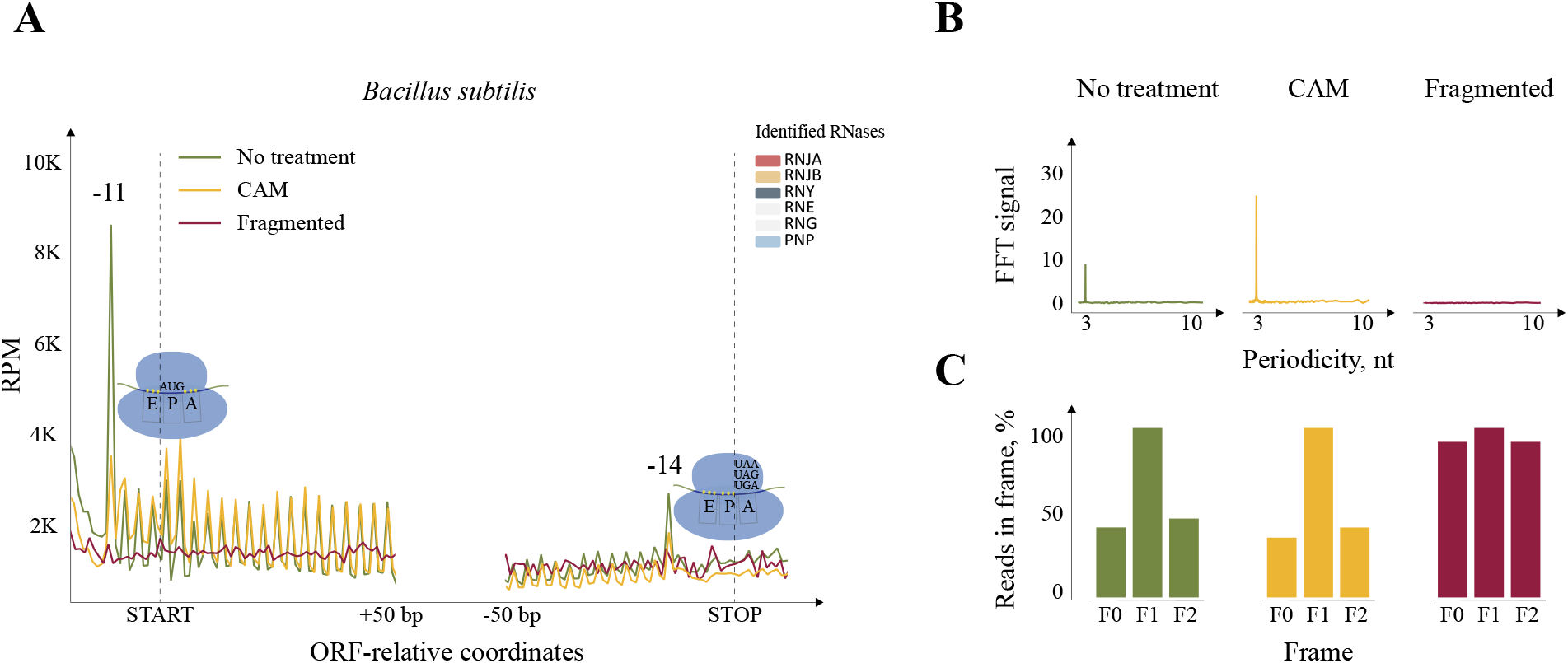
5’P mRNA sequencing can be used as a proxy for *in vivo* prokaryotic ribosome dynamics. **A**, Metacounts (5Pseq reads per Million) of 5’ mapping positions relative to translation initiation and termination codons of open reading frames (ORF’s) from *Bacillus subtilis*. (*19*). Exponentially growing cells (No treatment, in green), Chloramphenicol treated (CAM, in yellow) and randomly fragmented (in red) are shown. Identified RNases are highlighted in the upper right corner (unidentified are in light grey). Original 5PSeq positions are reported (no P-site correction was applied). **B**, Fast Fourier Transform (FFT) for the observed periodicity. **C**, Relative 5PSeq frame protection for all codons.

To assess the role of 5’-3’ exonculeolysis in shaping the co-translational mRNA degradation patterns across prokaryotes, we investigated other species with and without predicted 5’-3’ RNA exonucleases (Fig. S2) (*17*). We investigated representative species from the phylum Firmicutes (*Bacillus amyloloquefaciens, Lactobacillus plantarum, Lactobacillus reuteri*), Cyanobacteria (*Synechocystis sp PCC 6803*) and Proteobacteria (*Escherichia coli* and *Caulobacter crescentus*). *E. coli*, lacking 5’-3’ RNA exonuclease activity, generate only a subtle 3-nt periodicity (Fig. S2A), while a clear 3-nt pattern was observed in all other species (Fig. S2). In most cases we observed an accumulation of 5’P reads at the second nucleotide (F1) and at 11 and 14 nt upstream of start and stop codons (Fig. S2), as in *B. subtilis* (Fig. 1A). In *C. crescentus*, although we detected a clear 3-nt pattern, we did not observe footprints associated with slow ribosomes at the start or stop codons (Fig. S2B). Interestingly, in *Synechocystis*, the ribosome protection pattern was displaced by one extra nucleotide (*i.e*. accumulation at −12 nt from start and −15 nt from stop) (Fig. S2F). This could be explained by a different ribosome protection size, conformation or presence of potential cofactors(*8, 23*). Together these observations indicate that 5’-3’ exonucleolysis has a key role in shaping co-translational decay patterns, but also suggests the potential contribution of other factors in this process.

To confirm the causative role of the ribosome shaping the distribution of 5’P mRNA degradation intermediates, we perturbed ribosome dynamics using chloramphenicol (CAM), which is commonly used in prokaryotic ribosome profiling experiments(*22*). As expected for ribosomes stalled at elongation, we observed a clear increase of the 3-nt ribosome protection patterns in *B. subtilis* (Fig.1), *L. plantarum*, *L. reuteri* and *B. amyloliquefaciens* (Fig.S2). We also observed a relative excess of 5’P degradation intermediates in the 5’regions of ORFs and a decrease in the protection at the termination level, consistent with specific inhibition of translation elongation and not of initiation or termination(*22, 24*). This 5’P accumulation in the 5’regions of the ORFs was also evident in *E. coli* which displayed only a subtle 3-nt periodicity. All these demonstrate that ribosome protection commonly shapes the 5’P mRNA degradome in prokaryotes.

### 5’P serves as proxy for ribosome position and dynamics

Once demonstrated that 5’-3’ co-translational mRNA degradation is common among prokaryotes, we investigated up to what degree the abundance of 5’P mRNA decay intermediates can serve as a proxy for codon-specific pauses and ribosome dynamics. By aligning 5’P reads for each respective amino acid, we generated amino acid specific metagene profiles (Fig. 2 and Fig. S3). In *L. plantarum* this showed a clear 5’P accumulation associated with slow ribosomes at stop and Cysteine codons, likely associated with limited Cysteine in the growth media (Fig. 2A). We then perturbed the translation process and investigated its consequences on 5’P accumulation. First, we tested our ability to detect codon-specific pauses after 10 min treatment with Mupirocin (MUP). MUP is an antibiotic targeting isoleucyl t-RNA synthetase, and thus it is expected to stall ribosomes at Isoleucine codons(*22, 24*). As predicted, MUP treatment led to a clear accumulation of 5’P reads 14 nucleotides upstream of Isoleucine codons in *L. plantarum* and *L reuteri* (A-site stall, Fig. 2A and S3). This demonstrates a causal effect of the ribosome position shaping the prokaryotic degradome. Next, we investigated subtler codon-specific pauses associated with 5 minute CAM treatment in *B. subtilis*, *L. plantarum, L. reuteri and B. amyloliquefaciens* (Fig. 2A, B, Fig. S3). In addition to a general inhibition of translation elongation, CAM also leads to context-specific accumulation of ribosomes(*25, 26*). Using ribosome profiling it was previously shown in *E. coli* that CAM treatment led to artifactual ribosome footprints when Alanine (and less frequently Serine, Threonine or Glycine) are positioned in the E site. Reassuringly, 5PSeq is also able to discover CAM induced ribosome stalling 8 nt upstream of Alanine and Serine in the tested species (Fig. 2A, B and Fig. S3). This shows that the previously described context-specific CAM pauses are conserved across prokaryotes.

**Fig. 2.**
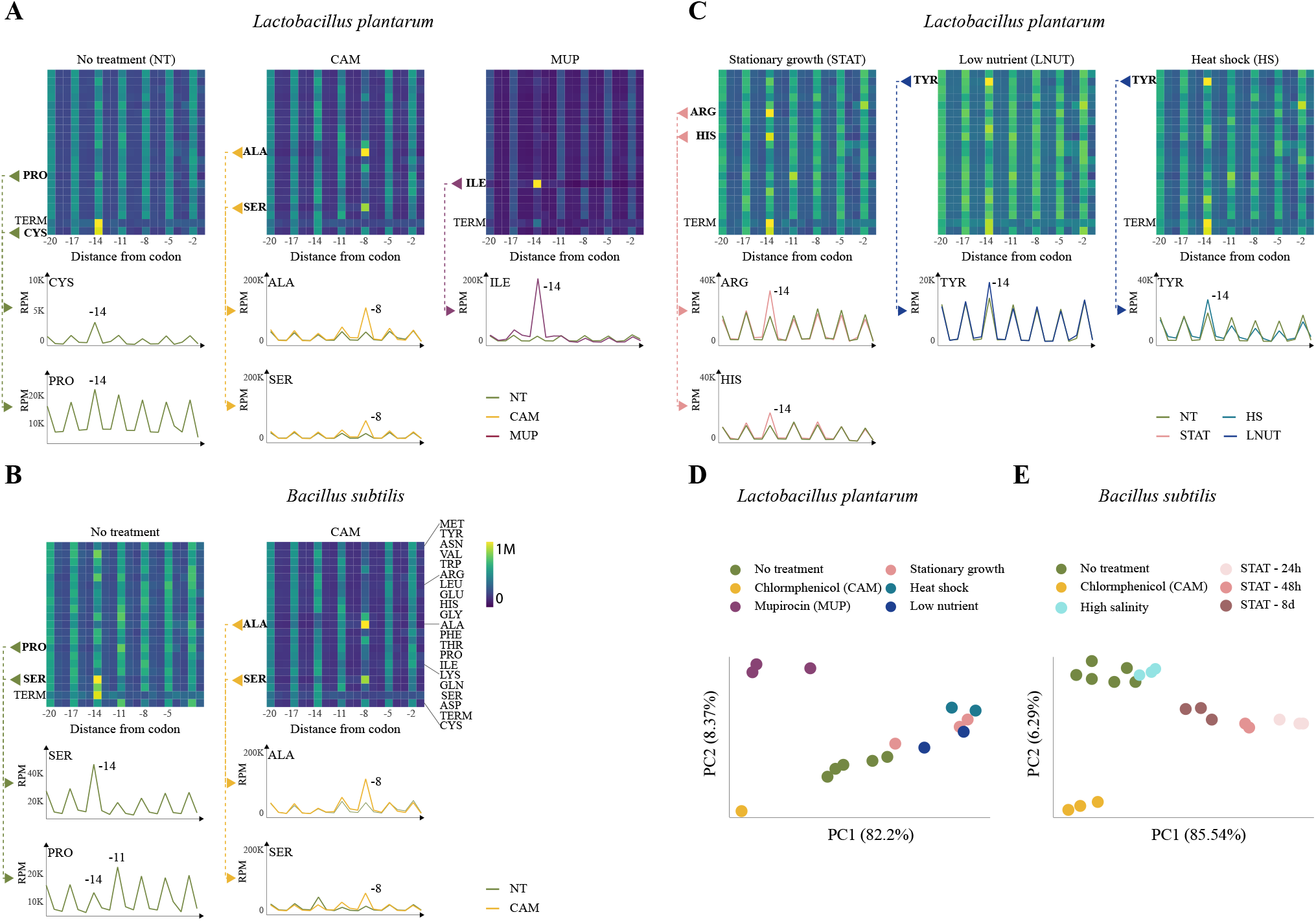
5PSeq reveals codon specific ribosome pausing patterns associated with stress and antibiotic treatment. Heatmaps for amino acid specific 5’P RNA coverage from blue (low) to yellow (high). Distance from specific amino acids is indicated as the number of nucleotides. Ribosomes paused at −14 indicates an A-site stall. Selected amino acids are shown also as line plots. **A,** *Lactobacillus plantarum* pauses after Chloramphenicol (CAM, yellow) and Mupirocin (MUP, purple) treatment. **B**, *Bacillus subtilis* context specific Chloramphenicol pauses induced by second-to-last amino acid (of peptide-chain) shown at the −8 position (CAM, yellow). **C**, *Lactobacillus plantarum* pauses after stress treatment. **D**, Principal component analysis of ribosome protection phenotype (see methods) allows distinguishing stress and drug treatment.

Next, we set out to investigate environmentally regulated changes in ribosome position. We studied the 5’P RNA degradation profiles under various stress conditions in *B. subtilis* and *L. plantarum* (Fig. 2C, Fig. S3). We identified clear codon pauses in *L. plantarum* at Tyrosine during heat shock and low nutrient conditions, and at Histidine and Arginine in stationary growth (Fig. 3C). As 5PSeq provides an easy overview of the ribosome-dependent accumulation of 5’P degradation intermediates at all amino acid positions, we used this information to generate a general amino acid-specific “ribosome protection phenotype” for each sample (see Materials and Methods). Using this phenotype and principal component analysis (PCA), we could clearly separate control, stressed and drug treated samples in *L. Plantarum* (Fig. 2D). The first two principal components also clearly separated cases of MUP and CAM treatments, stress conditions and time course measurements of stationary-phase *B. subtilis* (24h, 48h and 8d) (Fig. 2E). This showed that codon-specific ribosome protection phenotypes can be leveraged to distinguish across drugs and stress conditions. In addition to global patterns, 5PSeq also provides information regarding gene-specific ribosome stalls. We measured the relative strength and 3-nt protection patterns across genes (gene-specific protection index, see methods for details). This provides a measure of the relative stalling of the last translating ribosome (*i.e*. comparing peaks and valleys) independent of mRNA abundance for each gene. Drug and stress treatments induce changes in the protection index of individual genes involved in bacterial responses. For example, salt stress induced decreased pausing in genes involved in cell wall organization and iron-sulfur cluster assembly, in agreement with previous proteome-level studies (Table S3)(*27, 28*). All these demonstrate that 5PSeq can inform about ribosome position and translation regulation during stress, while detecting fast (*i.e*. in minutes) phenotypical changes in the translation process in response to drugs.

**Fig. 3.**
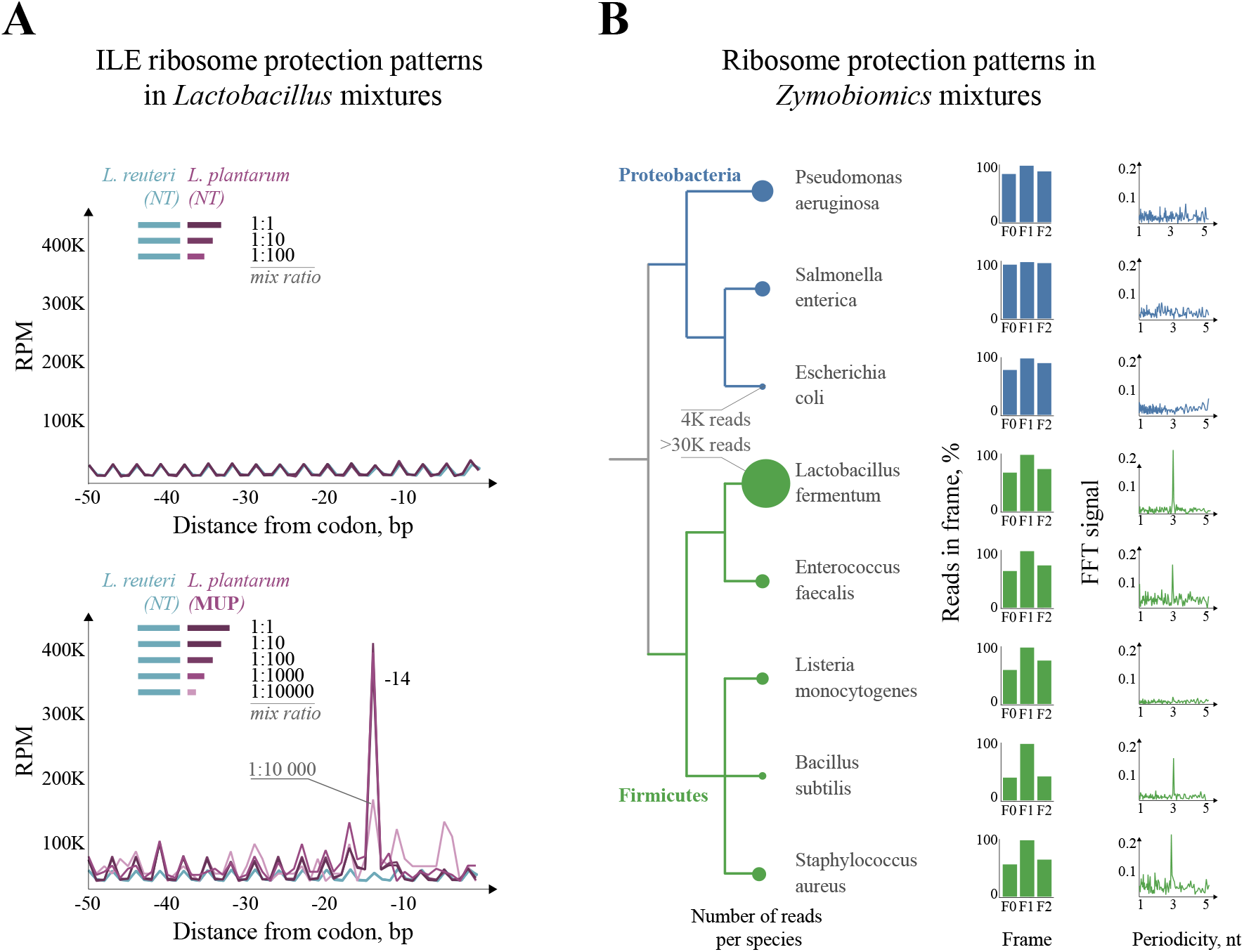
5PSeq enables the study of species- and codon-specific ribosome pauses in complex samples. **A,** Line plots showing 5PSeq metagene abundance in respect to Isioleucine (Ile) codons for *L. reuteri* (light blue) and *L. plantarum* (purple) mixed at different rations (from 1:1 to 1:10,000; *L. plantarum vs L. reuteri*). Mupirocin treatment of *L. plantarum* (MUP, bottom) leads to clear Ile pauses in respect to non-treated *L. reuteri* (NT). **B**, 5PSeq analysis from frozen cell suspension from the ZymoBIOMICS Microbial Community Standard (intended for DNA analysis). Numbers of assigned RNA reads are marked in circles. Relative frame protection and Fast Fourier Transform (FFT) as in Fig.1.

### 5PSeq enables the study of ribosome positioning in complex microbiome samples

Ribosome profiling, based on polyribosome fractionation followed by *in vitro* RNA footprinting and sequencing, is the current gold-standard for genomewide measurement of ribosome positions(*29*). Despite its multiple advantages, ribosome profiling is not well-suited to the study of complex mixtures of species. Ribosome profiling can only be applied to samples from which polyribosome fractions can be cleanly isolated (*i.e*. culturable microorganisms or tissue samples) and the ribosome protection fragments it produces are generally short, 23-24 nt(*22, 30*). In contrast, 5PSeq investigates longer, *in vivo* generated ribosome-protected fragments, which enables the distinction of closely related species within a complex sample (Fig. S4), does not require subcellular fractionation, and can be applied to RNA previously isolated and stored for years(*9, 19*). As this would allow the investigation of ribosome positions in currently inaccessible microbiome samples (metatranslatome), we set to demonstrate the feasibility of this approach.

We first investigated 5’P mRNA degradation intermediates in defined mixtures. We tested the ability of 5PSeq to detect species-specific perturbations in a mix of two closely related species, *L. reuteri* and *L. plantarum* (63-80% core nucleotide identity). We pooled different amounts of RNA from MUP-treated *L. plantarum* with untreated *L. reuteri* (from 1:1 to 1:10000, Fig. 3A). Reassuringly, the described MUP associated Isoleucine pause can be clearly detected even at the 1:10000 ratio (0.01% abundance). And this limit is simply driven by our sequencing depth (*i.e*. in the 1:10000 mix only 82 reads were mapped to *L. plantarum* coding sequences). Having confirmed our ability to detect species-specific perturbations, we investigated the ribosome dynamics in more complex cellular mixtures. We applied 5PSeq to RNA extracts of a defined microbial community standard for metagenomic analysis. We confirmed the existence of a clear 3-nt periodicity in *Lactobacillus fermentum, Enterococcus faecalis, Staphylococcus aureus, Listeria monocytogenes and Bacillus subtilis* from phylum *Firmicutes*, while it was absent in *Pseudomonas aeruginosa, Escherichia coli and Salmonella enterica* from phylum Proteobacteria (Fig. 3B). This demonstrates our ability to investigate ribosome dynamics in cells harvested for a different objective.

### Metatranslatome analysis of clinical and environmental microbiomes

Having proven that 5PSeq is suitable for analysis of complex synthetic mixtures, we decided to investigate clinical and environmental microbiome samples. We first studied the metatranslatome of vaginal microbiomes using previously isolated RNA samples. In addition to RNA-based indication of species abundance, we obtained clear 3-nt profiles for most identified species (Fig. 4A). This data allows for investigating *in vivo* ribosome protection phenotypes across species and patients. We can clearly observe sample clustering according to individual bacterial species, reflecting species-specific ribosome protection phenotypes in healthy hosts (Fig. 4B). In addition to global profiles, we can also investigate species-specific *in vivo* ribosome footprints (Fig. 4 and Supplementary Data 1). This opens the possibility to investigate species-specific ribosome dynamics in unculturable species, something that has not been possible until now.

**Fig. 4.**
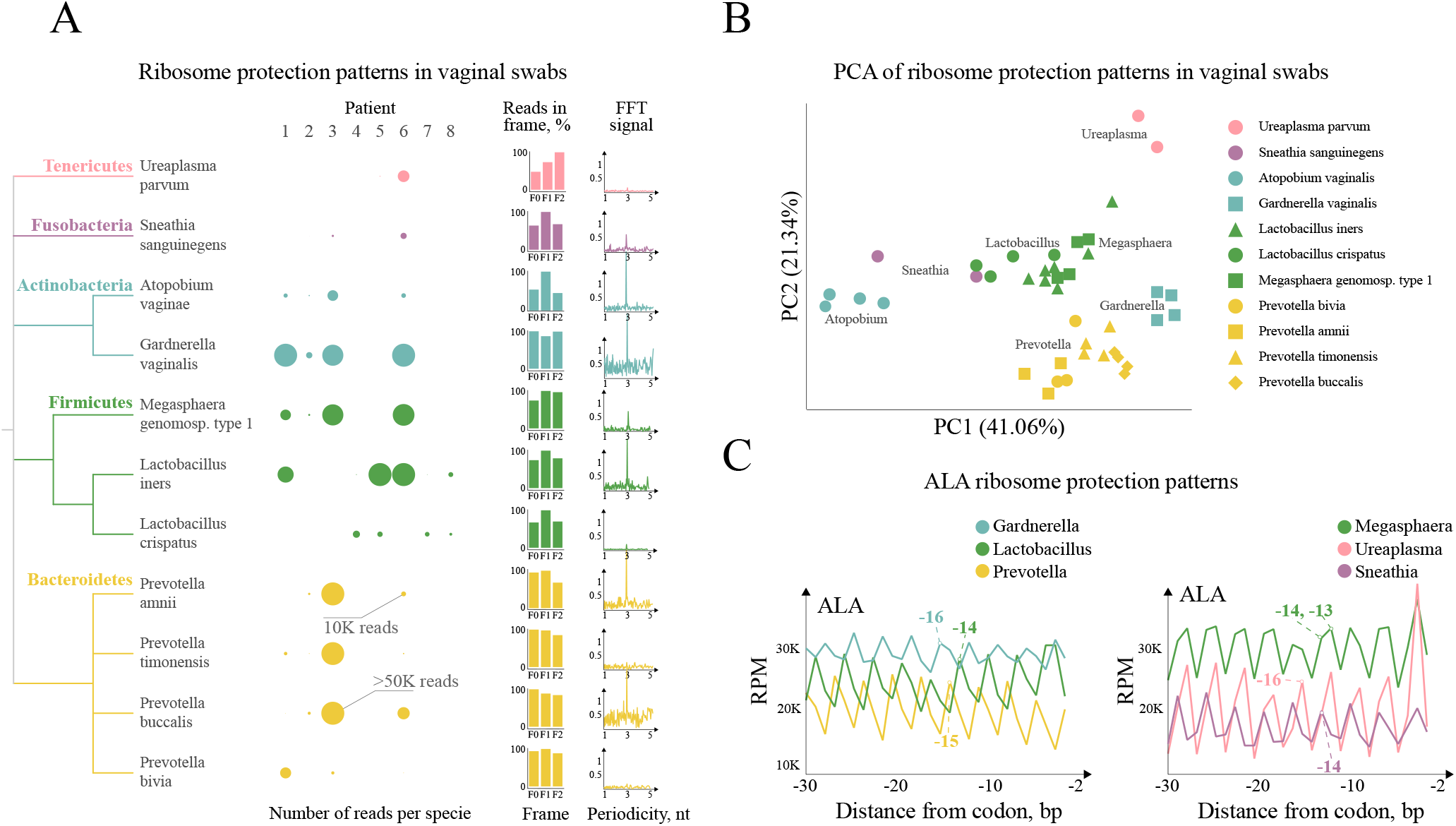
Ribosome dynamics in vaginal microbiome samples. **A,** 5PSeq analysis from previously isolated vaginal microbiomes. Number of assigned reads to each species and patient are marked in circles. Relative frame protection and Fast Fourier Transform (FFT) as in Fig.1. **B**, Principal component analysis of ribosome protection phenotype across clusters of phylogenetically close species. **C**, Example of *in vivo* amino acid specific ribosome protection in vaginal microbiomes.

Once demonstrated that 5PSeq can be applied to clinical microbiome samples, we expanded our study to more complex microbiomes(*31*). We investigated the human fecal microbiome and identified clear 3-nt patterns in multiple species (*e.g*. across Firmicutes and Bacteroidetes) (Fig. S5A). Finally, we applied 5PSeq to investigate translational phenotypes of bacteria found in compost (Fig S5B), revealing the applicability of our method to environmental microbiomes. Similar to the findings of the vaginal microbiome, species-specific ribosome protection phenotypes of feces and compost microbiomes show phyla-driven clustering (Fig S5C).

### 5PSeq can be used as a proxy for ribosome position across many prokaryotes

Having investigated the metatranslatome of both cultured species, clinical and environmental microbiomes, we used the generated data to investigate the conservation of co-translational mRNA decay across the prokaryotic tree of life (Fig. 5 and Fig. S6). In total we obtained information for 84 species with relatively high coverage distributed across 46 genera (Table S4 and S5). We can observe a clear 3-nt periodicity across most genera (inner bar plot in red). As previously described for *Synechocystis* (Fig. S2), different species present slightly different ribosome protection size. Protection in F1 (in green) is common across Firmicutes (e.g. *Lactobacillus, Bacillus, Megasphaera*), while protection in F2 (in yellow) is more common in Bacteroidetes (e.g. *Parabacteroides, Alistipes, Bacteroides*). This suggests that the size of the ribosome-protected footprints is similar across phylogenetically related species. Additionally, many species present more complex protection patterns. For example, *Ureaplasma* (Tenericutes), *Akkermansia* (Verucomicrobia) present a dual protection pattern in F1 and F2, while *Caulobacter* (Proteobacteria) and *Prevotella* (Bacteroidetes) in F1 and F0. However, it is important to note that ribosome-protected fragment size can also be influenced by environmental conditions or drug treatments(*8, 23*). Finally, as expected, a few species for which we do have good sequencing coverage (inner grey circle), such as *E. coli* (representative of class Gammaproteobacteria), do not present clear evidence of 5’P co-translational decay (Fig. 5).

**Fig. 5.**
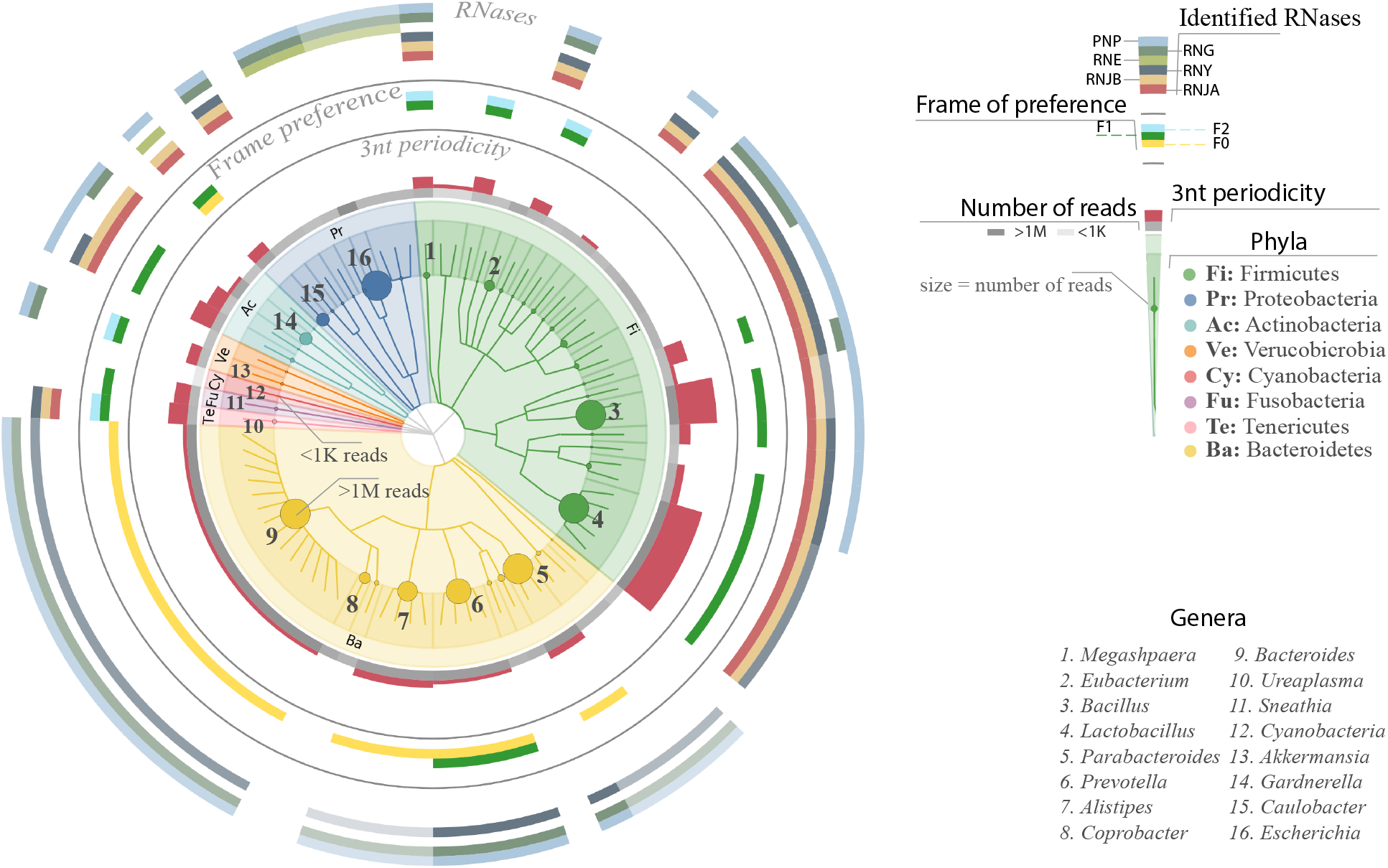
Co-translational mRNA decay is conserved across prokaryotes. From inside to outside: Taxonomic tree of investigated species; grey circle (number of assigned reads); red bars (strength of the 3-nt ribosome protection periodicity, protection frame preference (F0 yellow, F1 green, F2 light blue); Presence of selected enzymes involved in RNA degradation at genus level (RNJA (Ribonuclease J1), RNJB (Ribonuclease J2), RNY (Ribonuclease Y), RNE (Ribonuclease E), RNAG (Ribonuclease G) and PNP (Polyribonucleotide nucleotidyltransferase). Overall, 46 genera present in samples from cultured bacteria and complex environments, including a Zymobiomics mixture, vaginal swabs, feces and compost. In complex samples, species with at least 1000 reads in the coding regions (300 in case of compost) were considered and respective genera were analyzed. See methods for details.

Finally, we investigated the observed 5’P mRNA degradome profiles in relation to the phylogenetic conservation of the prokaryotic mRNA degradation machinery (outer circle in Fig. 5) focusing on species with >100K reads (Fig. S6) (*32, 33*).We observed agreement between the 3-nt periodicity and frame preference patterns with the taxonomic classification. Genera containing the 5-3’ exonuclease Ribonuclease J, (RNJA in red and RNJB in brown), such as *Bacillus*, *Lactobacillus* and *Caulobacter*, mostly present clear periodicities. While species more dependent on RNase E and G for mRNA decay (in light and dark green colors, respectively) such as *E. coli* (Gammaproteobacteria) present little 3-nt periodicity. Interestingly many species without conserved RNase J and RNase E (RNE), but with endonucleases RNase Y (RNY), RNase G (RNG) and 3-5’ exonuclease PNPase (PNP), like Bacteroidetes, also present clear 3-nt periodicity, albeit with a different ribosome protection pattern. All these suggest that in addition to the 5’-3’ RNA exonuclease activity, other prokaryotic nuclease activities are also able to shape the degradome in respect to the translation process.

## Discussion

Here we have shown that co-translation mRNA degradation is widespread among prokaryotes and that measurement of 5’P mRNA decay intermediates enables the study of ribosome dynamics. We have demonstrated the ability of 5PSeq to detect *in vivo* codon- and gene-specific ribosome dynamics without the need for drug treatment, subcellular fractionation or *in vitro* RNA degradation. We confirmed that it can easily detect environmentally triggered translational regulation in multiple species, both at codon- (Fig. 2) and gene-specific levels (Table S3). We also demonstrate that 5PSeq can easily detect fast (*e.g*. within 5 minutes) and specific perturbations of the translational process induced by drug treatments (Fig. 2). This opens new avenues to study environmental or chemical regulation of the translational process without the need for culturing or subcellular fractionation. Additionally, studying faster post-transcriptional regulation may help to disentangle direct from indirect secondary effects. We show that 5PSeq can be widely applied to complex clinical and environmental microbiomes to obtain species-specific metatranslatome profiles. By eliminating the need for complex experimental protocols and bacterial culturing, we can now investigate ribosome protection phenotypes in uncultivable bacterial species, which are estimated to comprise more than half of the human bacterial communities(*34*). More importantly, we can investigate it without the need for treatments that might alter cellular physiology or other biological processes. We show that our approach can detect translational responses even in species present at 0.01% abundance in a bacterial community, and by using longer ribosome footprint reads we can analyze complex bacterial communities at single-species resolution. 5PSeq enables studies of post-transcriptional gene expression regulation in microbiomes, and will complement current multi-*omics* approaches. As changes in ribosome position do not require variations in mRNA accumulation, 5PSeq is able to identify changes faster than transcriptomic approaches, as we describe for CAM. Complete understanding of all molecular mechanisms shaping co-translational mRNA degradation across the prokaryotic tree of life will require extensive mechanistic dissection. However, our results reveal that mRNA degradation pathways involving the exonuclease RNase J or the endonuclease RNase G (preferentially in the absence of RNase E) can shape 5’P mRNA degradation intermediates in respect to ribosome position (Fig. 5). We expect that in the future, metatranslatome analysis will enable the study of the unexplored post-transcriptional regulation in microbiome communities. This will bring new insights regarding how microbial communities interact with each other and their host, and how they respond to drugs and environmental challenges.

## Ethical matters

Collection and processing of sequenced vaginal samples was granted by the Regional Ethical Review Board in Stockholm (2017/725-31). Ethical approval for fecal samples was waived by the review board as only deidentified samples from healthy donors were used and no samples were stored in a biobank.

## Acknowledgements

We wish to thank members of the Pelechano, Kutter and Friedländer laboratories and to Ilaria Piazza for useful discussions. We kindly thank Sueli Marques, Deike Omnus and Jan Karlsen for assistance, Edmund Loh, Ciarán Condon, Kristina Jonas and Paul Hudson for generously sharing bacterial strains. Computational analysis was performed on resources provided by SNIC through Uppsala Multidisciplinary Center for Advanced Computational Science (UPPMAX). This project was funded by the Swedish Foundation’s Starting Grant (Ragnar Söderberg Foundation), the Swedish Research Council [VR 2016-01842], a Wallenberg Academy Fellowship [2016.0123], and Karolinska Institutet (SciLifeLab Fellowship, SFO and KI funds) to V.P.; the EU H2020-MSCA-IF-2018 program under grant agreement [845495 - TERMINATOR] to LN; the Söderbergs foundation to L.E.: Ministerio de Ciencia, Innovación y Universidades PGC2018-098073-A-I00 MCIU/AEI/FEDER, UE to J.H.C; the National Key R&D Program of China (2017YFC0908405) and National Natural Science Foundation of China (81870187) to W.W.; and an ERC Advanced investigator grant (742804) to L.M.S. VP and WW acknowledge the support from a Joint China-Sweden mobility grant from STINT (CH2018-7750) and the National Natural Science Foundation of China (81911530167) respectively.

## Author Contributions

VP, SH & LN conceived and designed the study. SH performed experimental work with support from DB & JW. LN and MR performed data analysis with support from JHC, SH & DB. WW & LMS contributed to initial conceptualization and data interpretation. LE contributed to data interpretation, design and supervision. SH, LN and VP drafted the initial manuscript and all authors revised it. VP supervised the study.

