## Supplementary material for "RNA degradation analysis reveals ribosome dynamics in complex microbiome samples"

#### Materials and methods

##### Bacterial growth and microbiome samples.

Overnight bacterial Cultures (not exceeding $OD_{600}$ 1) were used to dilute main culture to a starting $OD_{600}$ of 0.03-0.05. Culture was harvested by centrifugation when reached the logarithmic phase ($OD_{600}$ 0.4- 0.8) unless indicated otherwise. If not stated differently, bacterial cultures were grown at 37˚C and rotating, using the recommended growth Media.

In detail, *Lactobacillus plantarum* (ATCC 8014) and *Lactobacillus reuteri* (DSM 17938) were grown in MRS Broth (Sigma-Aldrich). *L. plantarum* stress treatments were carried out as follows: Stationary-phase cultures were grown for 27 hours post inoculation and harvested at $OD_{600}$ ~4.5. To generate samples for untreated control, heat shock and low nutrient, biological replicate cultures (40ml) were grown to mid-log phase ($OD_{600}$ 0.3-0.6), split (10ml for untreated control, 15ml for heat shock and 15ml of low nutrient sample) and cells were harvested by centrifugation. Untreated control pellet was flash frozen immediately after for RNA analysis. Prior to heat shock, cells were resuspended in prewarmed MRS broth and incubated in Thermomixer for 15 minutes at 60°C. Low nutrient cell pellets were washed excessively with 50ml 0.5xLB media, centrifuged and supernatant was completely removed. Cells were then resuspended in prewarmed 0.5xLB media (Sigma) and harvested after a total incubation time of 15 minutes at 37˚C.

*Bacillus subtilis* (strain 168 trpC2) was cultured in 2xYT (1.6% (wt/vol) Tryptone (Bacto), 1% (wt/vol) Yeast extract (Bacto) and 0.5% (wt/vol) NaCl) For extended growth, we collected samples at mid-log, 24 hours, 48 hours and eight days post inoculation. Salt stress was performed by mixing equal Volume of 2M NaCl with mid-log phase grown *B. subtilis* ($OD_{600}$ 0.5-0.6) followed by 10 minutes incubation at room temperature and harvest by centrifugation. *Bacillus amyloliquefaciens* and *Escherichia coli strain DH5α* (Invitrogen, Cat. No. 18265-017) were grown in LB Media. Biological Replicate Cultures of *L.plantarum* (ATCC 8014), *L.reuteri* (DSM 17938), *E.coli* (DH5å Invitrogen) and *Bacillus amyloliquefaciens* were grown to log phase and split to generate samples for untreated control, random fragmented control, Chloramphenicol(CAM) and Mupirocin (Mup) (only for *L.plantarum* (ATCC 8014) and *L.reuteri* (DSM 17938)). Chloramphenicol (CAM) was added to mid-log phase grown cultures at a final concentration of 100µg/ml, incubated for 5 minutes at 37˚C and subsequent harvested on ice containing additional 100µg/ml CAM (*9*). Mupirocin treatment (final 65µg/m MUP, Sigma-Aldrich) of mid-log *L. plantarum* and *L.reuteri* was carried out for 10 minutes at 37˚C following centrifugation and flash freezing of pellet. *Caulobacter crescentus* (strain NA1000) was grown in PYE Media containing 0.2% (wt/vol) peptone (Bacto), 0.1% yeast extract (Bacto) at 30˚C to mid-log, followed by centrifugation and flash freezing of cell pellet. *Synechocystis strain* PCC6083 was cultured in BG11 growth Media at 30˚C with a light intensity of 30 µE and 1% of atmospheric CO2 and harvested at mid-log phase(*35*).

##### RNA extraction

RNA was extracted (if not stated otherwise) as described in(*36*) with minor modifications In brief, cell pellets were resuspended in equal volume of LET (25 mM Tris pH 8.0, 100 mM LiCl, 20 mM EDTA) and water saturated Phenol pH 6.6 (Thermo Fisher). Cells were lysed with acid washed glass beads (Sigma-Aldrich) by vortexing for three minutes in MultiMixer. Following the addition of equal volumes of phenol/chloroform isoamyl alcohol pH 4.5 (25:24:1) and nuclease free water, lysis was extended by additional two-minute vortexing followed by centrifugation. Resulting aqueous phase was purified in two steps using phenol-chloroform isoamyl alcohol (25:24:1) followed by chloroform. After centrifugation, the clean aqueous phase was precipitated with sodium acetate-ethanol. For Lactobacillus mixtures (Fig3A), Microbial RNA extracted from *L.plantarum* (Untreated and Mupirocin treated) were mixed prior to RNA ligation step of 5PSeq Library protocol at different ratios with RNA extracted of *L.reuteri* (Untreated).

Technical replicates of the Microbial Community Standard (Zymobiomics Cat#D6300, Lot# ZRC 190633) consisting of eight deactivated bacterial strains, were generated by extracting RNA from 75-125µl thawed cell suspension.

Vaginal swab samples were mechanically lysed using beads in 1000µl of DNA/RNA shield (ZymoResearch) and lysate was stored at -80°C for 2 months before use. Lysate was thawed and 250µl was used to extract microbial RNA.

Feces from a healthy donor was collected and transported in 40% Glycerol. Technical replicates of RNA were extracted on the same day as stated earlier with minor modifications listed as follows. In brief, 500µl Feces-Glycerol suspension was mixed with equal volume of LET Buffer containing SDS (25 mM Tris pH 8.0, 100 mM LiCl, 20 mM EDTA, 10% SDS) and water saturated Phenol pH 6.6 (Thermo Fisher). Lyses was done by vortexing and carbide beads. Lyses duration was extended to 10 minutes after the addition of equal volumes of phenol/chloroform isoamyl alcohol pH 4.5 (25:24:1) and nuclease free water. All subsequent steps were performed as already indicated.

Compost RNA was extracted with 2g starting material (from Sundbyberg, Sweden) using RNeasy PowerSoil Total RNA Kit (Qiagen) as recommended in manufacturer's guidelines.

For all extractions, quality of RNA was assessed by either loading 1µg of total RNA on a 1.2% Agarose Gel or 12ng on a BioAnalyzer using an RNA Nano 6000 chip (Agilent Technologies).

**Polyribosome fractionation**

Was performed as described in Huch *et al*(*37*) with minor modifications. In brief, *B. subtilis* (168trpC2) was cultured in LB Media to mid-log phase at 37˚C and harvested on ice, containing 100µg/ml Chloramphenicol, by five-minute centrifugation. Resulting pellet was lysed in 1xTN (50mM TRIS/HCl pH 7.4,150mM NaCl, 1mM DTT, 100µg/ml CAM and Complete EDTA Free Protease inhibitor tablet) using glass beads and vortexing for 2 minutes, following a 5 min incubation on ice. Lysis and incubation procedure was repeated twice. Cell debris was cleared by centrifugation at 1500g for 5 minutes at 4˚C and supernatant was loaded on to a 15-50% sucrose gradient having an 80% cushion. After ultracentrifugation at 36,000 rpm for 90 minutes $Abs_{254}$ was monitored and fractionated. Subsequently, RNA was extracted from sucrose fractions by adding equal volumes of phenol/chloroform isoamyl alcohol pH 4.5 (25:24:1) and nuclease free water followed by two-minute vortexing and centrifugation. Aqueous phase was further cleaned up by the addition of Chloroform, vortexing and centrifugation. Resulting aqueous phase was sodium acetate-ethanol precipitated.

**Preparation of 5P Sequencing Libraries**

5PSeq libraries were prepared as previously described(*19*) with minor modifications using 150-9000ng total RNA as an input. To prepare random fragmented samples (negative controls), ribosomal RNA was depleted from DNA-free RNA and subsequent fragmented by incubating five minutes at 80˚C in fragmentation buffer (40mM Tris Acetate pH 8.1, 100mM KOAc and 30mM MgOAc). Reaction was purified using 2 volumes of RNACleanXP beads (Beckman Coulter) as recommended by the manufacturer. Free 5'OH sites were re-phosphorylated using 5 Units of T4 Polynucleotide kinase (PNK, NEB) and incubated at 37˚C for 60 minutes as recommended by the manufacturer. Re-phosphorylated fragmented RNA was purified using Phenol:Chloroform: Isoamyl Alcohol (24:25:1), followed by sodium acetate-ethanol precipitation. From this step forward, procedures for random fragmented and standard 5PSeq library preparation merge(*19*).

RNA was Ligated to either rP5_RND or rP5_RNA oligo (specified in Table S2) containing unique molecular identifiers. Ribosomal RNA was depleted using Ribozero rRNA removal kit (Illumina) suitable for Bacteria, Yeast and Human samples. Ribosomal RNA depleted sample was purified using 1.8V of Ampure beads (Abcam) and fragmented with heat (80˚C) for 5 min in 5x Fragmentation Buffer (200mM Tris Acetate pH 8.1,500mM KOAc, 150mM MgOAc). Subsequent samples were reverse transcribed using random hexamers to prime. Resulting cDNA was bound to streptavidin beads (M-280), subjected to enzymatic reactions of DNA end repair, fill-in of adenine to 5' protruding ends of DNA fragments using Klenow Fragment (NEB). Common adaptor (P7-MPX) was ligated and 5PSeq Libraries were amplified by PCR (15-17 cycles), purified using 1.8V of Ampure beads (Abcam) and quantified using Qubit (Thermo Fisher. Library size was estimated from bioanalyzer traces. 5PSeq Libraries were pooled by mixing equal amounts of each sample, following enrichment of 300-500nt size fragments.

HT-5PSeq Libraries were generated as recently described(*21*). In brief, DNA-free RNA was ligated with RNA oligos containing unique molecular identifiers. Ligated RNA was reverse transcribed priming with oligos containing a random hexamer and an Illumina compatible region. RNA was eliminated by addition of NaOH. Ribosomal RNA was depleted by adding in-house rRNA DNA oligo depletion mixes (Table S1) to the cDNA and performing a duplex-specific nuclease (DSN, Evrogen) treatment. rRNA depleted cDNA was PCR amplified (15-17 cycles). Depletion of ribosomal RNA with Ribozero Illumina (for bacteria and yeast) was done after the single-stranded RNA ligation step. Ribosomal depleted RNA was purified and reverse transcribed using the same oligos as stated above, and then amplified by PCR. Libraries were quantified by fluorescence (Qubit, Thermo Fisher), size estimated using an Agilent Bioanalyzer and sequence using a NextSeq500 Illumina sequencer.

**Data availability**

Sequencing data will be deposited on GEO. Clinical sample information will be deposited in dbGaP under access control.

**Sequence data pre-processing and mapping**

Demultiplexing and fastq generation of sequencing bcl image files was performed using bcl2fastq (version 2) with default options. Adapter and quality trimming was performed with the bbduk program of the BBTools suite (https://sourceforge.net/projects/bbmap/), with options {qtrim=r, ktrim=r, hdist=3, hdist2=2, K=20, mink=14, trimq=16, minlen=30, maq=16}, using BBTools default adapter set and polyG or polyA sequences for short reads. In order to reduce computational time, reads with both identical unique molecular identifier (UMI) and insert sequence were de-duplicated prior to mapping using the dedupe program of the BBTools suite with its default parameters. UMI sequences found in the first eight bases of each read were extracted using UMI-tools (version 1) with default options (using --bc-pattern NNNNNNNN).

Bacterial genomes were downloaded from the National Center for Biotechnology Information Assembly database (https://www.ncbi.nlm.nih.gov/assembly/) with the search terms: "bacteria"[Filter] AND (latest[filter] AND ("representative genome"[filter] OR "reference genome"[filter]) AND (all[filter] NOT "derived from surveillance project"[filter] AND all[filter] NOT anomalous[filter])) on March 21$^{st}$, 2019. The list was further filtered to include only one strain per species, giving priority to genomes marked as "reference". The resulting 5804 genomes (Table S4) were used to build the reference index. The index was built with the *bbmap* program of the BBTools suite, with default options (and k=10). Besides the reference index containing the 5804 bacterial genomes, separate indexes were built for individually cultured species and genus-level groups. The genomes for the latter groups were chosen from the initial set of 5804 genomes. Alignment was performed with the *bbmap* program of the BBTools suite, with the {32bit=t -da -eoom k=11 strictmaxindel=10 intronlen=0 t=16 trd=t minid=0.94 nzo=t} parameters. Alignment files were sorted and indexed with SAMtools (*38*)Deduplication based on UMIs was then performed with UMI-tools (version 1)(*39*) with options { --soft-clip-threshold 1 --edit-distance 2 --method unique}. The BAM files were then processed to count the number of reads in each species. We have used a pre-stored dictionary of chromosome and species names and used a custom script to perform counts in each species. Distribution of counts between genes coding for rRNA, tRNA, mRNA and other RNA types was computed with bedtools [https://bedtools.readthedocs.io/en/latest/ ]. The counts at mRNA coding genes were used to select top species in the complex samples as described below.

Individually cultured species were directly mapped to their reference indexes. All Zymobiomics mixtures, vaginal, fecal and compost microbiome samples were aligned to the bacterial reference index including 5804 species. We chose species with at least 1000 reads in the coding regions in all the samples except for compost, where we relaxed the selection to 300 reads, as there were less species with high counts. In total, 83 bacterial species with specified coverage belonging to 46 genera were identified in all the samples. Reference indexes were built for those 46 genera (species were chosen from the preselected list of 5804), and all the complex fecal and compost samples were separately aligned to those references.

**Fivepseq and ribosome dynamics analysis**

Deduplicated alignment files, along with genome sequence and annotation files, were provided as input to our recently developed *fivepseq* package (*20*) for analysis and visualization of 5' endpoint distribution of reads with default options applied. *Fivepseq* provides information regarding presence of 3-nt periodicity (FFT, Fast Fourier transform), distribution of 5' counts relative to CDS start/stop or to nucleotides within each codon (translational frames), and codon and amino acid specific protection patterns. *Fivepseq* analyzes only one genome per run. Thus, alignment files for complex samples were used as input for fivepseq for each genome separately. For genus-level analysis, sequence and annotation files for individual species were concatenated into one.

To generate a "ribosome protection phenotype" we took the sum of counts positioned 30 to 1 nucleotide upstream of each amino acid and concatenated the per amino acid scaled counts to obtain a vector that describes ribosome protection in each sample. These vectors were used as input for principal component analysis (PCA) performed with the *prcomp* function of the R package *stats* (v 3.6.1). The PCA plots were generated with the *autoplotly* package (v0.1.2) in R.

In addition to the global pattern of ribosome pausing in either of the nucleotides in each codon (F0, F1 or F2), we have also computed gene-specific changes in the protection index upon different treatments. For this, we considered the counts in each reading frame, excluding the first two and the last codon of each transcript. A likelihood ratio test of independence was used to estimate differences in frame preference in each gene in untreated and treated conditions. The p-values were derived from $X^2$ distribution of the log-likelihood statistic. Multiple testing correction was performed with Benjamini-Hochberg procedure (implemented in statsmodels package **v**0.11.1)(*40*). The effect size was reported using Cramer's V metric, adjusted for directionality of the change (Supplementary data 1). The adjusted effect size metric was used as input for gene set enrichment analysis (via *Webgestalt* (http://www.webgestalt.org/) along with reference functional annotation set obtained from the Uniprot database (https://www.uniprot.org/).

**Taxonomic analysis**

Taxonomic trees were generated with the *graphlan* tool (v0.9.7) (https://huttenhower.sph.harvard.edu/graphlan). The taxonomic lineage information for all the 84 bacterial species identified in our samples was downloaded from the NCBI Taxonomy database with the *efetch* program from NCBI *e-utilities*. Trees were annotated with information about the library size, 3-nt periodicity, preferred ribosome protection frame and presence of enzyme annotations for each genus (Supplementary data 1). The library size equaled the maximum number of mRNA reads per species per sample, brought to the range of 0 to 1 (≥1M reads) reads.

The 3-nt periodicity was computed taking into account the absolute value of Fast Fourier transform (FFT) signal for the 3-nt periodicity wave and the preference for ribosome protection frame, as computed by the *fivepseq* package. For FFT, the maximum of the signals for transcripts aligned either at the start or at the end was taken. The preference for ribosome protection frame was assessed based on the value of frame protection index (FPI), computed by the *fivepseq* package as $2{*}F_i/(\sum_{j=0}^{2} F_j - F_i)$ , for each frame $F_i$. The frame with maximum absolute FPI value was regarded as (mis)preferred, and the significance of the preference was assessed based on t-test p value comparing counts in the given frame with the other two combined (the FPI and p values are found in the *frame_stats.txt* file of the *fivepseq* output). A positive FPI value means that one of the nucleotides in each codon on average has higher counts (is preferred), while a negative FPI value means that one of the nucleotides on average has low counts (is misprefered), while the other two nucleotides receive more counts. *E.g.* if $F_1$ is preferred, it will have positive FPI value and will be highlighted in the tree as a single preferred frame of protection, while if say $F_2$ is (mis)preferred (has a negative FPI value), the tree will highlight $F_0$ and $F_1$ as the frames of preference. The FFT and FPI values were brought to the range of 0 to 1, and the maximum of the two values was taken to describe the strength of 3nt-periodicity.

Enzyme annotations were obtained from the EggNOG database (v5.0)(*32*).The presence of each enzyme in each genus was counted as a number between 0 and 1, depending on the fraction of species within the genus annotated with the enzyme. The tree highlights these values with corresponding opacity.

### Supplementary figures

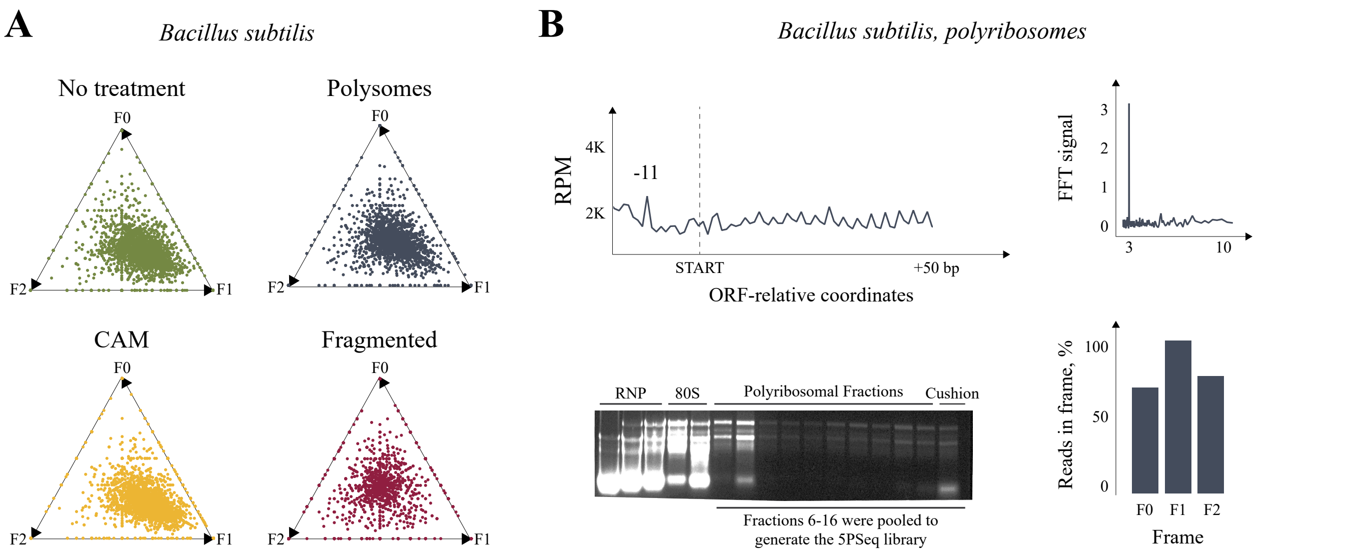


**Fig. S1. 5'P 3-nt periodicity is associated with co-translation mRNA decay.** a, Gene specific 3-nt protection for *Bacillus subtilis* as reported by *fivepseq*10. Each point corresponds to a gene and the proximity to the triangle boundaries (F0,F1,F2) their relative protection. Non-treated exponentially growing cells are shown in green, polyribosome associated mRNA degradation intermediates in blue, chloramphenicol treated cells in yellow and random fragmented (in red). b, 5PSeq metagene analysis of *Bacillus subtilis* after polyribosome fraction isolation (As in Fig.1). Bar on the bottom of the agarose gel indicates fractions used for 5PSeq library generation.

Figure S2

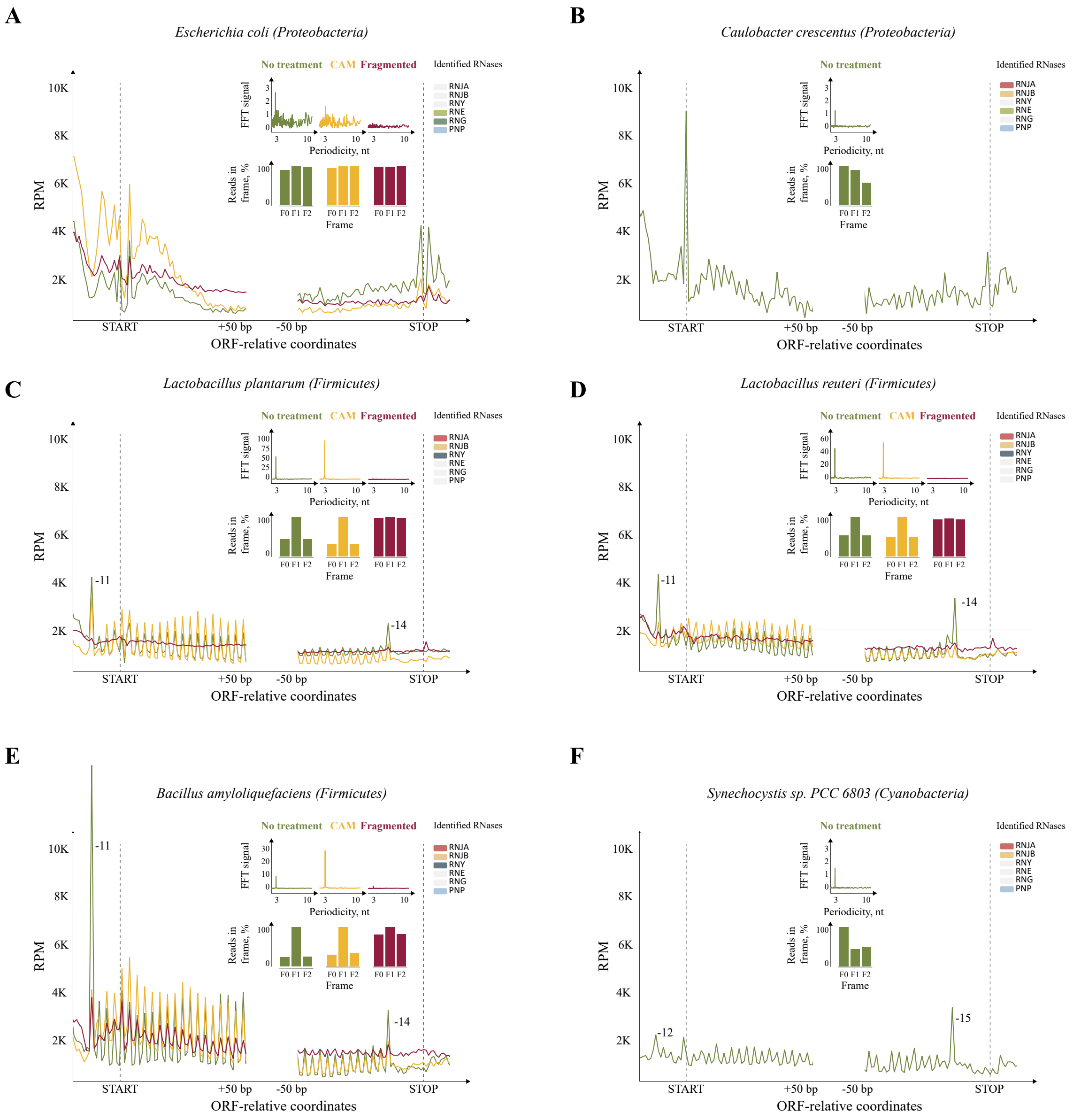


**Fig. S2. Ribosome associated 3-nt periodicity can be found in multiple prokaryotic species.** Metagene analysis for multiple species displaying metagene 5PSeq protection, Fast Fourier Transform (FFT), relative frame protection and identified RNases (as in Fig.1). a, *Escherichia coli*. b, *Caulobacter crescentus*.c, *Lactobacillus plantarum*. d, *Lactobacillus reuteri*. e, *Bacillus amyloliquefaciens* f, *Synechocystis sp.* PCC 6083.

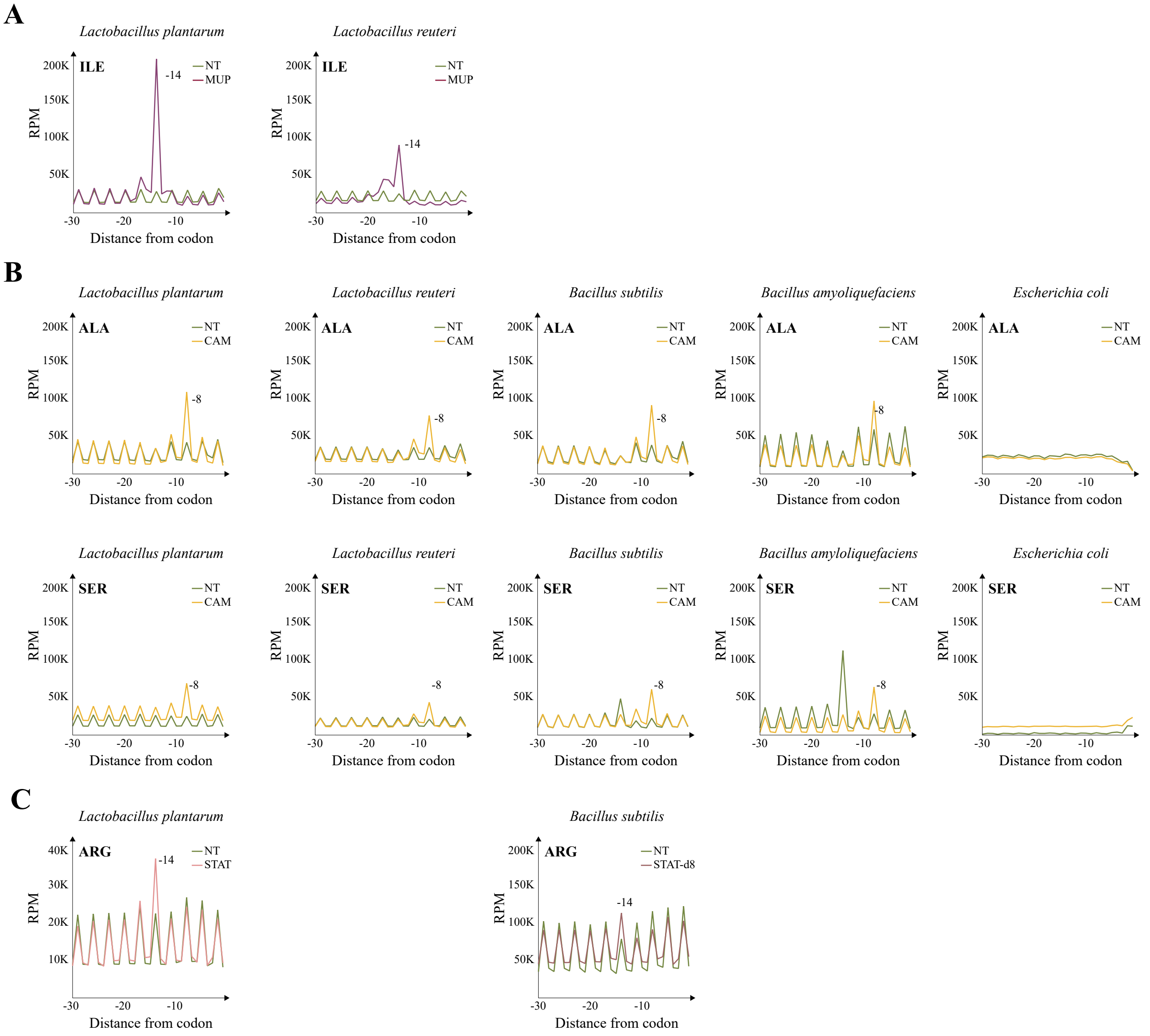


**Fig. S3. Species- and codon-specific ribosome pauses in response to stress or antibiotic treatment.** Line plots showing amino acid specific ribosome pauses as measured by 5PSeq. a, Isoleucine (Ile) pause comparing no treatment (NT) and mupirocin treatment (MUP) in *L. plantarum* and *L. reuteri*. b, relative Alanine (Ala) and Serine (Ser) ribosome pause as measured by 5PSeq for multiple species in response to chloramphenicol treatment (CAM). c, relative Arginine (Arg) pauses in stationary phase growth for *L. plantarum* (27 hours) and *B. subtilis* (8 days).

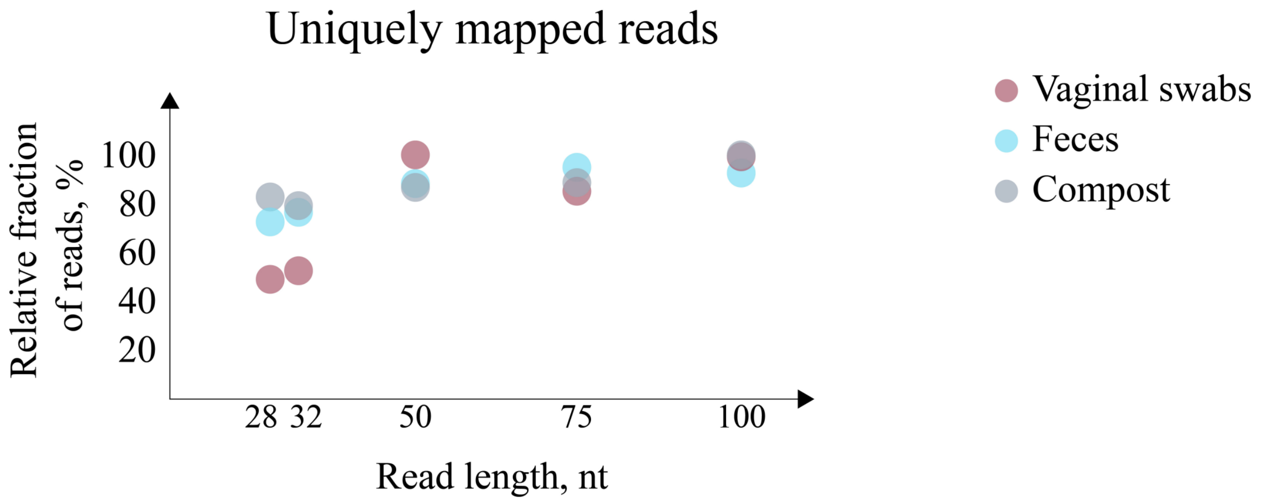


**Fig. S4. Longer ribosome protected 5PSeq reads enables better species specific assignment in complex microbiomes.** Shown are the relative number of uniquely mapped reads (scaled within each sample) as a function of read length.

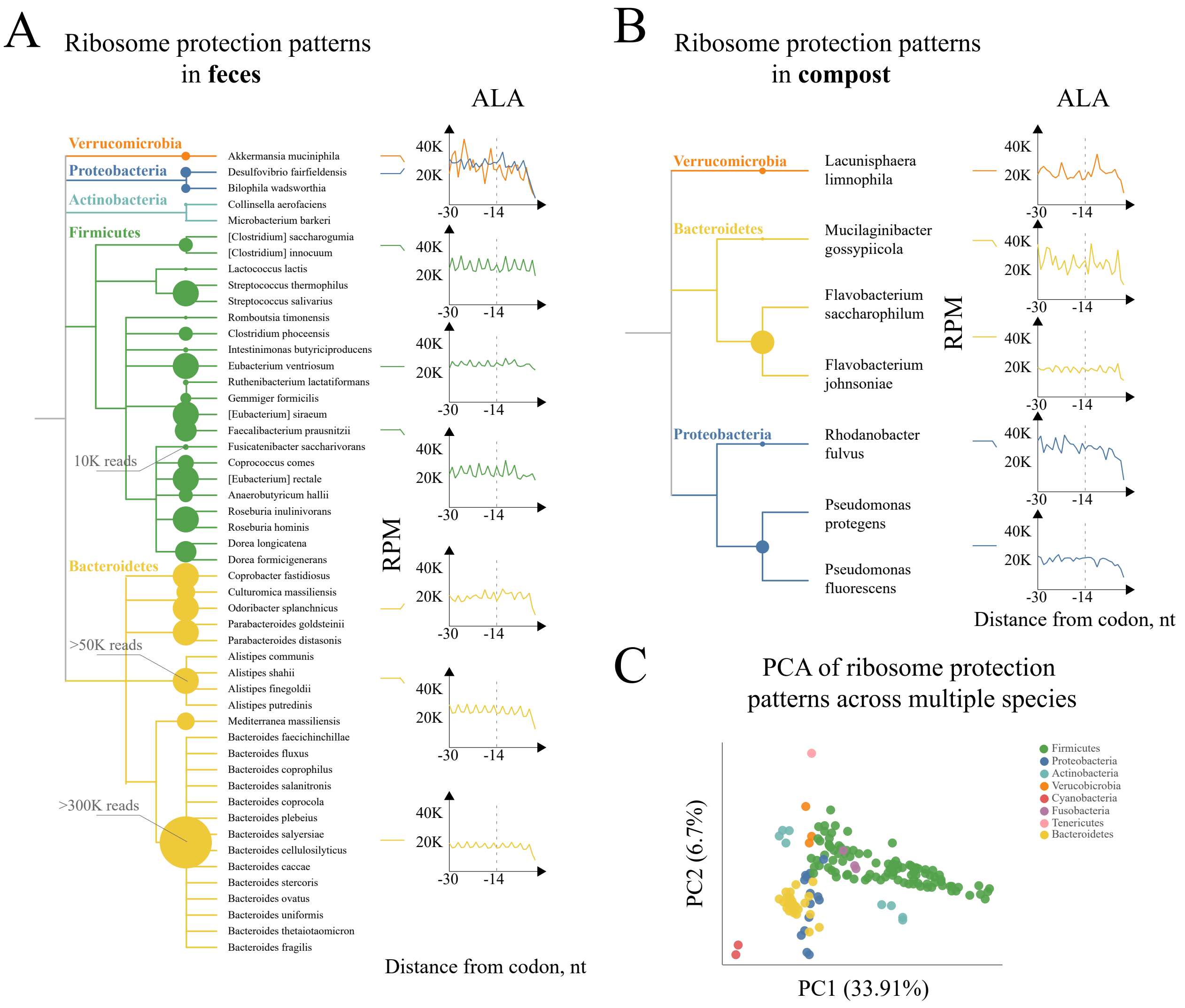


**Fig. S5. Ribosome dynamics in fecal and compost microbiomes. a,** 5PSeq analysis from fecal microbiomes. Number of assigned reads are marked in circles. Example of *in vivo* amino acid specific (Alanine) 3-nt ribosome protection periodicity for selected species. b, same for compost microbiome. **c**, Principal component analysis of ribosome protection phenotype across analyzed phyla (all the phyla from cultured and complex microbiome samples including 46 genera, with treatments).

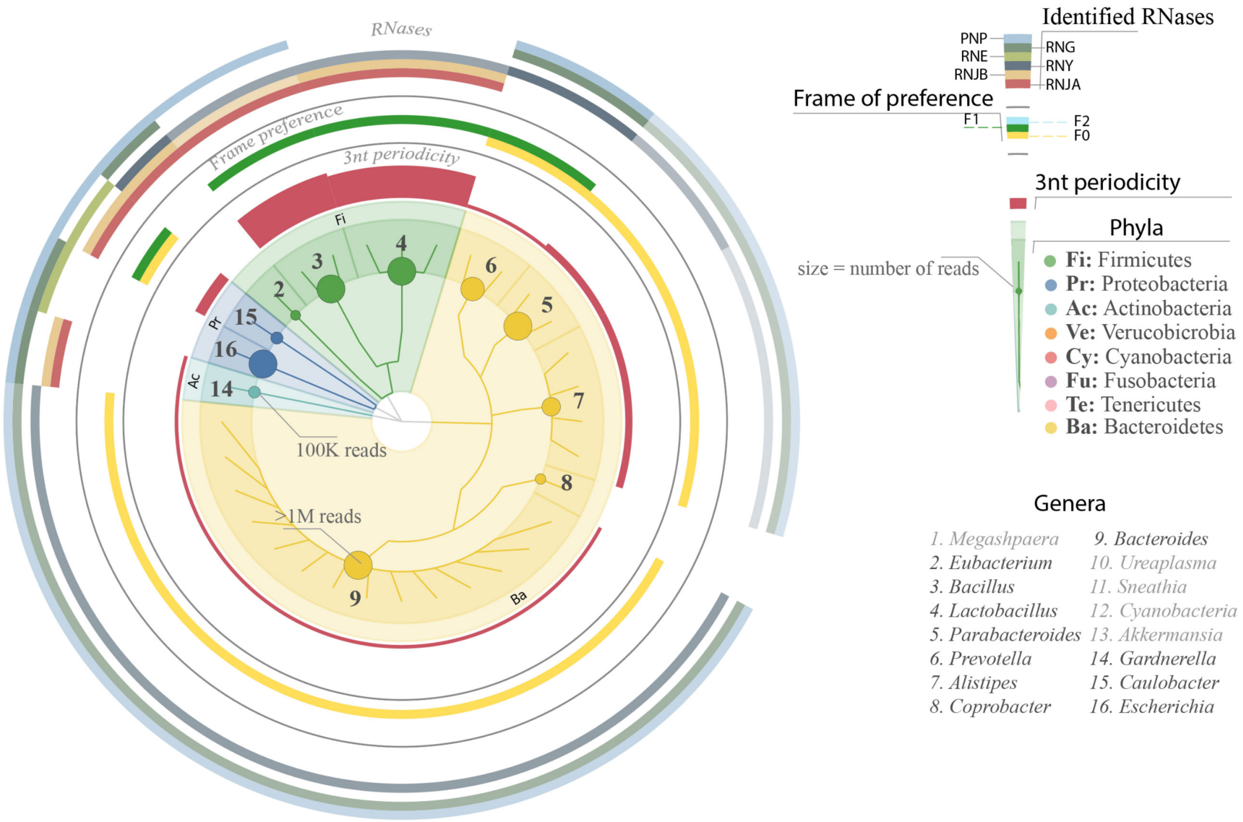


**Fig. S6. Co-translational mRNA decay in species with relatively high coverage.** From inside to outside: Taxonomic tree of investigated species; red bars (strength of 3-nt ribosome protection periodicity, protection frame preference (F0 yellow, F1 green, F2 light blue); Presence of selected enzymes involved on RNA degradation at genus level (RNJA (Ribonuclease J1), RNJB (Ribonuclease J2), RNY (Ribonuclease Y), RNE (Ribonuclease E), RNAG (Ribonuclease G) and PNP (Polyribonucleotide nucleotidyltransferase)), with opacity indicating the fraction of species with identified enzyme in each genus. Overall, 11 genera including species with at least 100K reads in the coding regions, from samples of cultured bacteria and complex environments, including a Zymobiomics mixture, vaginal swabs, feces and compost are analyzed. See methods for details.
